# Tryptophol acetate and tyrosol acetate, small molecule metabolites identified in a probiotic mixture, inhibit hyperinflammation

**DOI:** 10.1101/2021.12.16.472991

**Authors:** Orit Malka, Ravit Malishev, Marina Bersudsky, Manikandan Rajendran, Mathumathi Krishnamohan, Jakeer Shaik, Daniel A. Chamovitz, Evgeni Tikhonov, Eliya Sultan, Omry Koren, Ron N. Apte, Benyamin Rosental, Elena Voronov, Raz Jelinek

**Author notes:** These authors contributed equally to this work (Orit Malka, Ravit Malishev).

## Abstract

Probiotic fermented foods are perceived as contributing to human health, however solid evidence for their presumptive therapeutic systemic benefits is generally lacking. Here we report that tryptophol acetate and tyrosol acetate, small molecule metabolites secreted by the probiotic milk-fermented yeast *Kluyveromyces marxianus* inhibit hyperinflammation (e.g., “cytokine storm”). Comprehensive *in vivo* and *in vitro* analyses, employing LPS-induced hyperinflammation models, reveal dramatic effects of the molecules, added in tandem, on mice morbidity, laboratory parameters, and mortality. Specifically, we observed attenuated levels of the pro-inflammatory cytokines IL-6, IL-1α, IL-1β and TNF-α, and reduced reactive oxygen species. Importantly, tryptophol acetate and tyrosol acetate did not completely suppress pro-inflammatory cytokine generation, rather brought their concentrations back to baseline levels thus maintaining core immune functions, including phagocytosis. The anti-inflammatory effects of tryptophol acetate and tyrosol acetate were mediated through downregulation of TLR4, IL-1R, and TNFR signaling pathways and increased A20 expression, leading to NF-kB inhibition. Overall, this work illuminates phenomenological and molecular details underscoring anti-inflammatory properties of small molecules identified in a probiotic mixture, pointing to potential therapeutic avenues against severe inflammation.

## Introduction

Sepsis is a life-threatening condition manifested by severe inflammation leading to multiple organ dysfunction. The inflammatory host response associated with both innate and adaptive immunity mechanisms play an important role in the development of the clinical and pathological manifestations of sepsis [1]. Recently, it has been found that the prognosis of the disease is dependent not only on the virulence of the microorganisms, but, mainly, on the host response affected by the pathogen-associated molecular patterns (PAMPs) [2,3] response, often leading to multi-organ failure and adverse clinical outcomes with high mortality rates [4]. Hyperinflammation or severe inflammation occur in various disease conditions, including sepsis and septic shock [5] and acute stages in chronic diseases [6]. Severe inflammation was also shown to be a major cause of mortality from COVID-19 [7,8].

Despite the significant progress in inflammation treatment following the discovery of antibiotics, high mortality from sepsis still exists. Thus, new approaches to improve conventional therapies are highly sought. Varied food products have been touted to endow anti-inflammatory properties and as such attract significant interest [9]. Food-extracted substances have been particularly explored as anti-inflammatory agents, including probiotics [10,11], curcumin [12], resveratrol [13], plant extracts [14], and phenolic compounds from natural sources [15]. However, the therapeutic benefits of most such substances against severe inflammation have been limited [16].

Probiotics, particularly milk-fermented microorganism mixtures (yogurt, kefir), have been known to bolster the innate immune system and host-defense mechanisms against pathogens [17,18]. In a recent study, we reported on a yet-unrecognized mechanism for cross-kingdom inhibition of pathogenic bacterial communication and virulence by a small molecule - tryptophol acetate – secreted by the probiotic yeast *Kluyveromyces marxianus* in a milk-fermented probiotic microorganism mixture [19]. Specifically, tryptophol acetate was found to disrupt biofilm formation and reduce virulence of several human pathogenic bacteria, underscoring a novel mechanism for combating bacterial colonization and pathogenicity.

Here, we report that tryptophol acetate and tyrosol acetate, another *K. marxianus* – secreted metabolite, exhibit remarkable anti-inflammation activities, in *in vitro, ex vivo* and *in vivo* models. Through application of LPS-induced hyperinflammation, we observed that the two molecules had anti-oxidation, anti-inflammatory, clinical, histological and hematological systemic protective effects against severe inflammation. Importantly, tryptophol acetate and tyrosol acetate did not give rise to immune system shutdown, rather reduced pro-inflammatory cytokine production to baseline levels while retaining core immune processes including phagocytosis and generation of anti-inflammatory cytokines. We found that the anti-inflammatory activities of tryptophol acetate and tyrosol acetate are mediated through downregulation of *TLR4, IL-1R*, and *TNFR* signaling pathways and suppression of NF-κB activity. In particular, we observed that the molecules enhanced expression of A20, a key modulator of NF-κB signaling pathways [20]. Overall, this study demonstrates significant anti-inflammatory properties of probiotic yeast-secreted metabolites, underscoring their important therapeutic potential.

## Materials and Methods

### Materials

Dulbecco’s Modified Eagle Medium (DMEM), Roswell Park Memorial Institute (RPMI 1640), heat-inactivated fetal bovine serum (FBS) were purchased from Biological Industries (Beit Haemek, Israel). Penicillin, streptomycin, ECL western blotting substrate, opti-MEM, and Bradford reagent were purchased from Thermo-fisher. Dihydrochloride (AAPH), 2,2-Diphenyl-1-picrylhydrazyl (DPPH), 2,20–azobis (2-amidinopropane, fluorescein, hydrogen peroxide, sodium citrate, acetonitrile, formic acid, sulfuric acid, ethanol, methanol, sodium carbonate, lipopolysaccharide (from E. coli O55:B5, L2880), propidium iodide (PI) (Solution P4864), NaCl, Triton X-100, BSA, NaF, Na3VO4, AEBSF, polyethylenemine (PEI), Leupeptin, and Aprotinin were purchased from Sigma-Aldrich. Xylene was purchased from Epredia (lot 6601). Tris buffer was purchased from Bio-lab (Jerusalem, Israel). Thioglycolate was purchased from life technologies, 2085262 (ultra-pure grade). All reagents and solvents were of analytical grade and were used as received.

### Animal studies

#### C57BL/6 mice

All experiments were performed in accordance with the Institutional Animal Care and Use Committee at Ben-Gurion University (IL-43-07-2020). Eight-week-old C57BL/6 mice (Harlen, Rehovot, Israel) without a specific gender, weighing approximately 20 gr were held in specific pathogen-free conditions at the University Central Research Facility for at least one week prior to commencing studies. The mice were fed (*ad libitum*) with V1154-703 ssniff™ (Soest, Germany) and were allowed water continuously. Environmental enrichment was provided to all animals based on the requirements of the particular mouse strain. Temperature was maintained at 21°C, and animals were exposed to a 12 h light, 12 h dark cycle with a 15 min ramp-up and ramp-down to simulate dusk and dawn. The mice were divided into two study groups: untreated (mice injected with LPS and administered a placebo), Treated (mice injected with LPS and treated by oral administration of mixture of tryptophol acetate and tyrosol acetate). Mice were housed in groups of six per cage. LPS dose was 30 mg/kg to each mouse (~20 g), which was injected intraperitoneally.

### Treatment with tryptophol acetate and tyrosol acetate

Tryptophol acetate and tyrosol acetate were synthesized as previously described [21] and dissolved in DDW to desired concentrations (final concentration of 150 μg/kg, this concentration was chosen after dose response examination; data not shown). The molecular mixture was administrated through oral gavage in a volume of 200 μL (the maximum liquid volume that can be given to a mouse) twice per day, in the morning (08:30–09:30) and evening (19:30–20:30). DDW was used as a vehicle and was administrated orally to the control group at the same time.

In one set of experiments, the mixture of molecules was administrated concomitantly with LPS injection and treatment was continued two times per day for 3 days (72 hours). In another set of experiments, the first treatment with the molecules started 28 hours after LPS injection and was continued two times per day until the end time of the experiment (156 h).

### Clinical evaluation of inflammation

Clinical signs of inflammation were recorded twice daily. Measurement of body weight, signs of diarrhea and appearance of blood traces in stool, as well as signs of dyspnea were assessed. The mean percentage of weight loss was calculated daily as the ratio of measured body weight and weight measured before injection of LPS. Every 6 h after LPS injection till endpoint of the experiments (till 72 h), mice were anesthetized with 5% isoflurane (2-chloro-2-(difluoromethoxy)-1,1,1-trifluoro-ethane) and blood samples were got from tail vien. Lungs, livers, colons, and intestinals were obtained at the same time intervals and were cleaned from contents by flushing with 10 ml of sterile PBS. The tissues were taken for extraction of RNA and part of them were embedded in paraffin following a standard protocol. 5-μm sections using a rotary microtome (MRC-Lab, HIS-202A). Paraffin embedded tissues were cut into 5-μm sections using a rotary microtome (MRC-Lab, HIS-202A), rehydrated and stained with hematoxylin.

### Immunohistochemistry Staining

Tissue sections were de-paraffinized in xylene and re-hydrated with decreasing concentrations of alcohol. Subsequently, endogenous peroxide was blocked with hydrogen peroxide, and antigen retrieval was achieved by treating sections with 0.01M sodium citrate, pH 6.0 for 1 min in a pressure cooker. After blocking with universal blocking solution (ZYMED Laboratories, San Francisco, CA), tissue sections were stained with the designated primary antibodies. A Vectastain Elite ABC Peroxidase kit or Universal ImmPRESS kit (Vector Laboratories, Burlingame, CA) were used for secondary antibodies, and visualization was performed using 3-amino-9-ethylcarbazole (AEC) as a substrate (ZYMED Laboratories, San Francisco, CA). Sections were then stained with hematoxylin for counterstaining and mounted using VectaMount AQ Aqueous Mounting Medium (Cat-No. H-5501; Vector Laboratories). Myeloperoxidase (MPO)-positive (Abcam), F4/80-positive (Santa Cruz, 377009), and LY6C-positive cells (Abcam, ab15627) were counted in stained sections in six randomly chosen fields (×200), and bars are indicated standard errors of the means.

### Blood count

Blood samples were collected from mice tails with EDTA micropipette capillaries (Exigo) and analyzed directly after collection by a veterinary hematology analyzer (Exigo H400, Boule Medical AB, Spånga, Sweden).

### Cytokine measurements by ELISA

To measure the extent of the inflammatory response in mice, serum obtained from the blood of LPS-treated and untreated mice, we customized a highly sensitive milliplex^®^ MAP Kit (Cat # MPXMCYTO-70K; Millipore) with color-coded beads and fluorescent dyes according to the manufacturer’s recommendations.

### *In vitro* macrophage experiments

#### 264.7 cell line

The RAW 264.7 cell lines (third passage) were purchased from American Type Culture Collection (ATCC1 TIB-71™). Cells were cultured in DMEM with 10% FBS and 1% penicillin/streptomycin, in an atmosphere of 5% CO_2_ and 95%humidity at 37°C. Cells were regularly tested for mycoplasma contamination.

### Peritoneal macrophages

Mouse peritoneal macrophages were obtained from C57BL/6J mice (6–8weeks old) 72 hours after intra-peritoneum injection of Thioglycolate and cultured in complete RPMI medium with and without LPS (100ng/ml) (O55:B5, L2880, Sigma) and in the presence or absence of the tryptophol acetate and tyrosol acetate (1+2 mixture).

### Cytokine production in peritoneal macrophages

Cells (3×10^5^) were seeded in 24-well plates and after 24 h, supernatants were collected and assessed for cytokine secretion. To measure the extent of the inflammatory response in the LPS stimulated peritoneal macrophages, we used murine ELISA kits according to the manufacturers’ recommendations for IL-1β (PeproTech, 900-K47), IL-6 (Biotest, DY406), TNF-α (Biotest, DY410), TGF-β (Biotest, DY1679), IL-1α (Biotest, MAB400-500; AB-monoclonal BAF400, AB-polyclonal 400-ML-005), and INF-β (BioLegend B292426).

### ROS analysis

Peritoneal macrophage cells were seeded (1× 10^5^ cells) into 96-well plates for ROS and phagocytosis analysis.

After 2 h of incubation in complete medium at 37°C, cells were stimulated with LPS (100ng\ml) and incubated overnight in the absence or presence of tryptophol acetate and tyrosol acetate (100 μM each). After 24 hours, the cells were washed two times with PBS and incubated for 1 h with CellROX Deep Red Reagent (Invitrogen, Carlsbad, CA) prior to analysis by flow cytometry. Dead cells were gated out using PI. The presence of ROS was analyzed by the fluorescence geometric mean (GM) of the CellROX Deep Red fluorescence.

### Phagocytosis analysis

Peritoneal macrophage cells were seeded in 96-well plates with RPMI medium. After 2 adherences at 37°C, all the cells were stimulated with LPS (100 ng\ml) and incubated overnight in the absence or presence of the tryptophol acetate and cytosol acetate mixture (100 μM each). After 24 hours the cells were washed two times with PBS and incubated for 1 h or 3 h at 37°C in 200 μl RPMI medium, alone (control) or with Fluoresbrite^®^ Yellow-Green carboxylate microspheres (1 μm) (Polysciences Inc., 15702) in a ratio of 3:1 beads-to-cells. In the flow cytometry analysis, dead cells were gated out using PI. Free beads were analyzed alone and gated out from the analysis. The fraction of live cells positive for beads was used to estimate phagocytosis functionality.

### Western blot analysis for NF-κB expression

Protein expression was monitored using Western blot analysis. 10×10^6^ murine peritoneal macrophages were seeded on petri dishes. After overnight incubation at 37°C the cells were stimulated with LPS (100 ng/ml) in the absence or presence of the mixture of tryptophol acetate and tyrosol acetate (100 μM each molecule) and incubated for 30 minutes and harvested in lysis buffer (50 mL 3M NaCl, 25 mL 1M Tris pH 7.5, 10 mL Triton X-100, and 10 mL 0.5M EDTA were mixed and added to 905 mL distilled water) in the presence of protease inhibitor cocktail (1:50 complete, Sigma-Aldrich, Israel). Lysates were placed on ice for 30 min and then centrifuged for 30 min (12,000 rpm) at 4°C. Supernatants (cytosolic fractions) were collected, and protein concentrations were determined using a Bio-Rad protein assay kit (lot 5000202, Israel). SDS sample buffer was added, and samples were boiled for 5 min and then frozen at −20°C until use. NF-κB (after 30 minutes) was assayed using cell lysates (70μg), separated on SDS-PAGE and analyzed using the following: antibody raised against the anti-NF-κB (p-65) (1:1000; abcam, ab32536), β-actin (1:1000; MP Biomedicals, Santa Ana, CA).

### Real-time PCR

Total mRNA from obtained tissues and RAW 264.7 macrophages was extracted using a RNA extraction kit (ISOLET II RNA Mini Kit, Bioline). cDNA was synthesized from 1 μg of RNA using a PrimeScriptTM RT reagent kit (Lifegene, Bio-52073). Subsequent real-time PCR was performed with an iCycler iQ™ Real-Time PCR Detection System (Bio-RAD). PCR results were analyzed with SDS 2.02 software (Applied Biosystems, Thermo). The level of target gene expression was calculated following normalization of the GAPDH gene level in each sample and presented as relative units. Quantitative PCR was performed with Taqman Master Mix (Rhenium, Israel) for: TLR4 (Cat 4453320, Assay ID Mm00445273_m1), Tnfrsf1b (Cat 4453320, Assay ID Mm00441889_m1), GAPDH (Cat 4453320, Assay ID Mm99999915_g1), and IL-1R (Cat 4453320, Assay ID Mm00434237_m1) or with Mix SYBR Green Master Mix (Applied Biosystems).

**Table 1.**
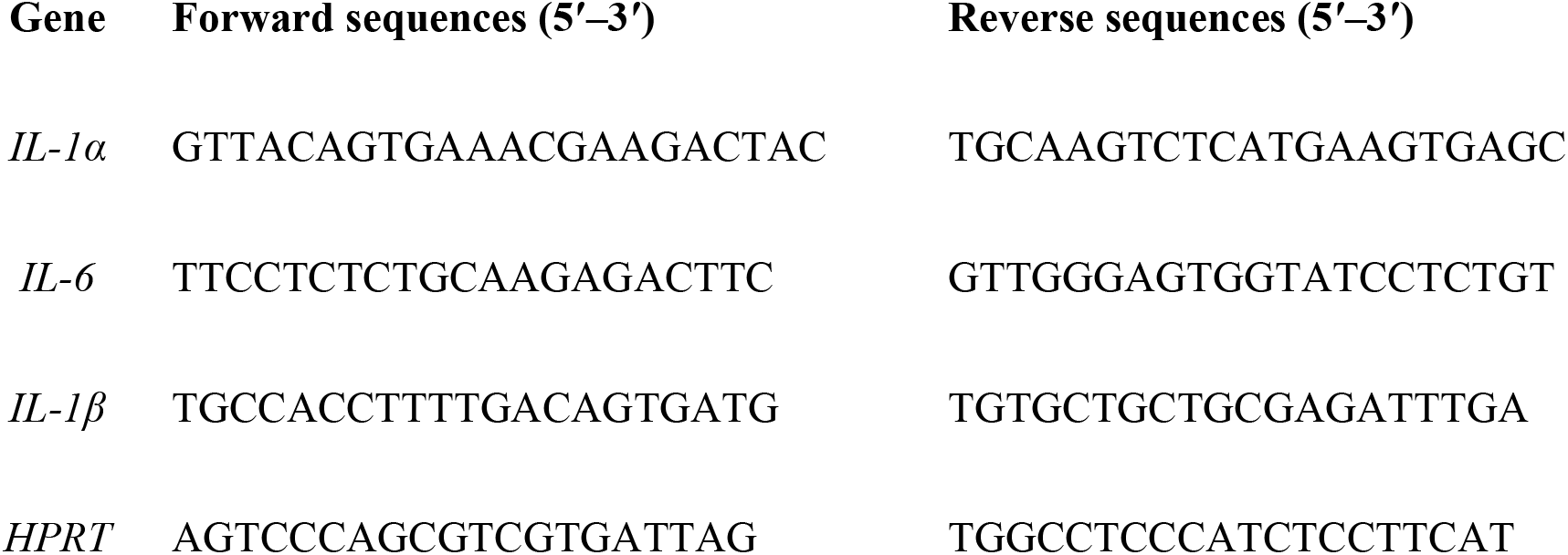
Primers used for RT-qPCR of target gene

### A20 expression in HEK-293 T cells

#### Cells

HEK-293 T cells were grown and maintained in Dulbecco’s Modified Eagle’s Medium (DMEM) at 37 °C and 5% CO_2_, supplemented with 10% fetal calf serum and antibiotics (50 units/ml penicillin, 50 μg/ml streptomycin).

### Antibodies and plasmids

Mouse monoclonal antibodies (mAbs) specific for A20 (Santa Cruz Biotechnology, Inc., Santa Cruz, CA). Mouse anti-β-actin mAbs (Merck Millipore, Darmstadt, Germany). Horseradish peroxidase (HRP)-conjugated goat anti-mouse Abs (Abcam Biotechnology). The full-length human or mouse A20 cDNA with a human influenza hemagglutinin (HA) tag cloned in a pCAGGA mammalian expression vector (HA-A20) was kindly provided by Dr. Shai Cohen (Cancer and Vascular Biology Research Center, The Rappaport Faculty of Medicine and Research Institute, Technion, Israel), previously described [22].

### Transient transfection in HEK-293 T cells

HEK-293 T cells were chosen for transient plasmid transfection. Briefly, 5 x 10^5^ HEK-293 T cells were plated and left to grow for 24 hours in the incubator (37 °C and 5% CO_2_). The following day, the old DMEM medium was replaced with serum free-fresh DMEM and the cells were left in the incubator for 30 minutes. DNA (plasmids) samples were prepared for transfection; 1.5 μg of purified HA-A20 plasmid DNA was added to 250 μL of Opti-MEM medium, then 5 μL of (final concentration of 5 μg/mL) of polyethylenimine (PEI) was added to the DNA, vortexed and incubated at RT for 20 minutes. The PEI-DNA mix was added to the cells steadily drop by drop and the cells were left in the incubator (37 °C and 5% CO_2_) for 4 hours. After the transfection, the cells were incubated with the Molecule mix (200 μM each) for time range of 1 and 2 h. Cells were harvested at indicated time points and lysed in lysis buffer containing 25mM Tris HCl, pH 7.5, 150mM NaCl, 5mM EDTA, 1mM Na_3_VO_4_, 50mM NaF, 10μg/ml each of leupeptin and aprotinin, 2mM AEBSF and 1% Triton X-100) followed by 30 min incubation on ice. Lysates were centrifuged at 13,000 rpm for 30 min at 4°C and the nuclear-free supernatants were used for further analysis. 20 μg of protein was resolved on 10% SDS-PAGE and blotted on to nitrocellulose membrane. Membranes were immunoblotted with mouse anti-A20 (1:2000) and mouse anti-β-actin mAbs (1:500) as indicated and visualized following ECL exposure.

### Protein concentrations

Protein concentration was measured by the method of Bradford using bovine serum albumin as standard.

### Electrophoresis and immunoblotting

Whole cell lysates were resolved by electrophoresis on 10 % polyacrylamide gels using Bio-Rad Mini-PROTEAN II cells. Gel proteins were electroblotted onto nitrocellulose membranes (Schleicher and Schuell) at 100 V for 1 h, using BioRad Mini Trans-Blot transfer cells. After 1 h of membrane blocking with 3% BSA in TBS at 37 °C, the membranes were incubated with the indicated primary Abs followed by extensive washings in TBST and incubation with HRP-conjugated goat anti-mouse. Immunoreactive protein bands were visualized using the enhanced chemiluminescent.

### Data analysis

All results are expressed as the mean ± SD or mean ±SE, as indicated. Data for all in-vivo and in-vitro experiments were analyzed by a one/two-way ANOVA followed by Tukey post-hoc test using GraphPad Prism (GraphPad Software, San Diego, Ca). A p-value of ≤ 0.05 was considered statistically significant.

### Fecal microbiome analysis

Total DNA was extracted from fecal samples using a PureLink microbiome DNA extraction kit (Invitrogen, Carlsbad, CA) followed by a 2-min bead-beating step, as previously described by Shouval, R. et al. [23]. Purified DNA was PCR amplified using PrimeSTAR Max (Takara-Clontech, Shiga, Japan) for the variable V4 region (using 515F-806R barcoded primers) of the 16S rRNA gene. Amplicons were purified using Agencourt AMPure XP magnetic beads (Beckman-Coulter, Brea, CA), and subsequently quantified using a Quant-It Picogreen dsDNA quantitation kit (Invitrogen, Carlsbad, CA). Equimolar DNA amounts from individual samples were pooled and sequenced using the Illumina MiSeq platform at the Genomic Center, Azrieli Faculty of Medicine, Bar-Ilan University, Israel. Sequencing data were processed using QIIME2 version 2019.10. [24]. Single end sequences with a similarity ≥ 99% were assigned as the same feature. Taxonomy was assigned using the GreenGenes database [25]. Chimeric sequences were removed with DADA2 (--p-trunc-len 160) [26]. Samples with <1000 features and features with total frequency <2 were filtered out. Rarefaction was done using 5,000 sequences per sample. Alpha and beta diversity were calculated based on rarefied datasets.

### Statistical analysis of microbiome results

The distribution pattern of quantitative variables was examined using a Shapiro-Wilk test. Quantitative variables were compared between groups with a Wilcoxon U-test, for non-normally distributed data, respectively. All tests were 2-tailed, and in all, p ≤ 0.05 was considered significant. Bacterial alpha diversity was assessed by the Shannon index. Statistical significance of microbial alpha diversity differences was confirmed using a Kruskal–Wallis test, followed by paired Mann-Whitney tests, with Benjamini-Hochberg correction for the false discovery rate. Beta diversity was calculated using weighted UniFrac distances. Statistical significance was confirmed using a permutational multivariate analysis of variance (PERMANOVA). Differences in relative abundances of bacterial taxa between groups were identified using the ANCOM method [24].

## Results

Scheme 1 depicts the chemical structures of tryptophol acetate (**1**) and tyrosol acetate (**2**), recently identified as metabolites secreted by the probiotic fungus *Kluyveromyces marxianus* in milk-fermented microorganism mixture (“kefir”), and also shown to exhibit intriguing antibacterial properties through blocking quorum sensing [21]. Tryptophol acetate and tyrosol acetate also exhibit significant anti-oxidative properties when tested together (Figure 1, SI); this feature was an important factor in examining their anti-inflammatory properties, depicted herein. The experiments outlined below were carried out using synthetic tryptophl acetate and tyrosol acetate (99% purity).

**Fig. 1.**
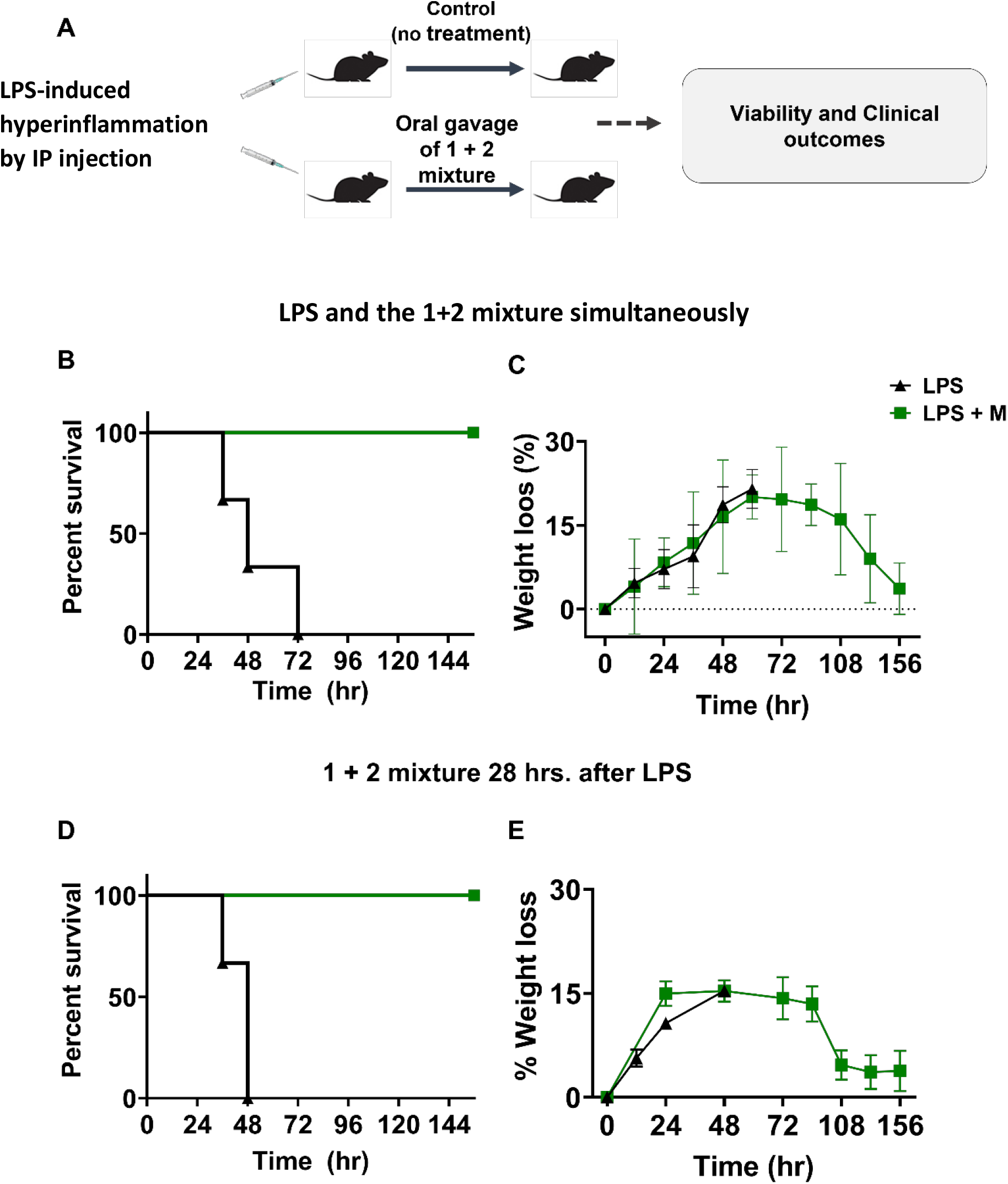
Tryptophol acetate and tyrosol acetate mixture shield mice from LPS-induced severe inflammation. **A.** Experimental scheme. After intraperitoneal (IP) injection of LPS (30 mg/kg), mice were randomly divided into two groups, PBS-treated and treated with **1**+**2** immediately after LPS injection. **B - C**. A third group treated with the molecules 28 hours after LPS injection (**D, E**). Survival (**B, D**) and weight loss (**C, E**) of the mice were monitored for 156 hours.(n=6 in each group)

**Scheme 1.**
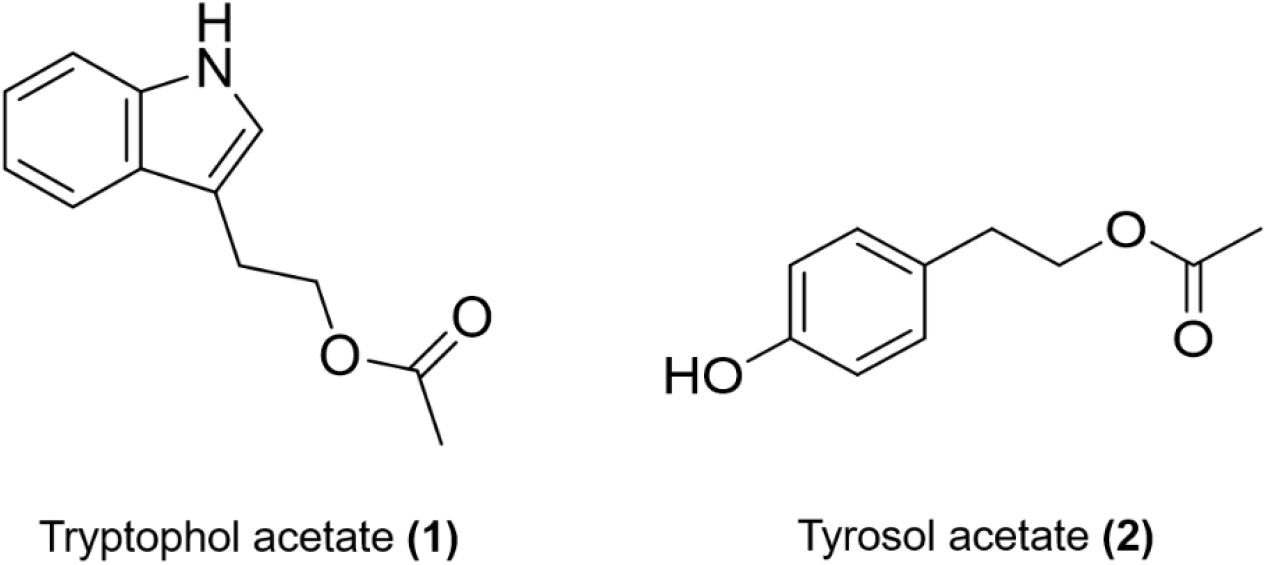
Chemical structures of tryptophol acetate (1) and tyrosol acetate (2).

### Oral uptake of tryptophol acetate and tyrosol acetate mixture provides protection against LPS-induced cytokine storm in mice

We investigated the anti-inflammatory properties of the **1+2** mixture (1:1 mole ratio) *in vivo* using a model of lipopolysaccharide (LPS)-induced hyperinflammation, which triggers a massive pro-inflammatory cytokine release that occurs in sepsis [27,28]. C57BL/6 mice were injected with LPS (30 mg/kg), and the effects of the **1** + **2** mixture administrated orally by gavage (each molecule at 150 μg/Kg per mouse) were monitored until the mice regained their initial weight (156 hours) (Figure 1A).

Initially, mice were injected with LPS and at the same time interval were treated orally with **1+2** mixture. The percentage survival and weight loss in the LPS-administered mice, with and without treatment, show that the molecules mixture appreciably improved the disease outcome. Specifically, while all LPS-treated mice died within 72 hours due to severe inflammation, 100% of the mice orally given the mixture of **1** + **2** survived (Figure 1B). Furthermore, the LPS-administered mice experienced substantial weight loss prior to mortality (Figure 1C, black line). In contrast, mice that were orally treated with the molecular mixture initially lost weight (up to 60 hours) but reverted to their initial values within 156 hours (Figure 1C, green line).

To test the therapeutic effects of the molecules, we carried out another experiment, in which **1+2** mixture was given to mice that were already impacted by the onset of LPS-induced hyperinflammation (Figure 1D-E). In these experiments, the molecular mixture was administered to the mice by gavage 28 hours after LPS injection. At this time-interval, severe signs of the disease were already evident, including decreased motor activities, ruffled fur, diarrhea, substantial eye discharge and respiratory distress. The results in Figure 1D demonstrate that while all untreated mice died within 48 hours after LPS injection, 100% of the mice treated with the **1** + **2** mixture survived. The weight lost patterns in mice treated with the molecules 28 hours after LPS administration when hyperinflammation was already developed (Figure 1E) were similar to the case of mice that were orally given the two molecules simultaneously with LPS (e.g., Figure 1C). Overall, the results presented in Figure 2 demonstrate a pronounced therapeutic effect of the tryptophol acetate and tyrosol acetate mixture.

**Fig. 2.**
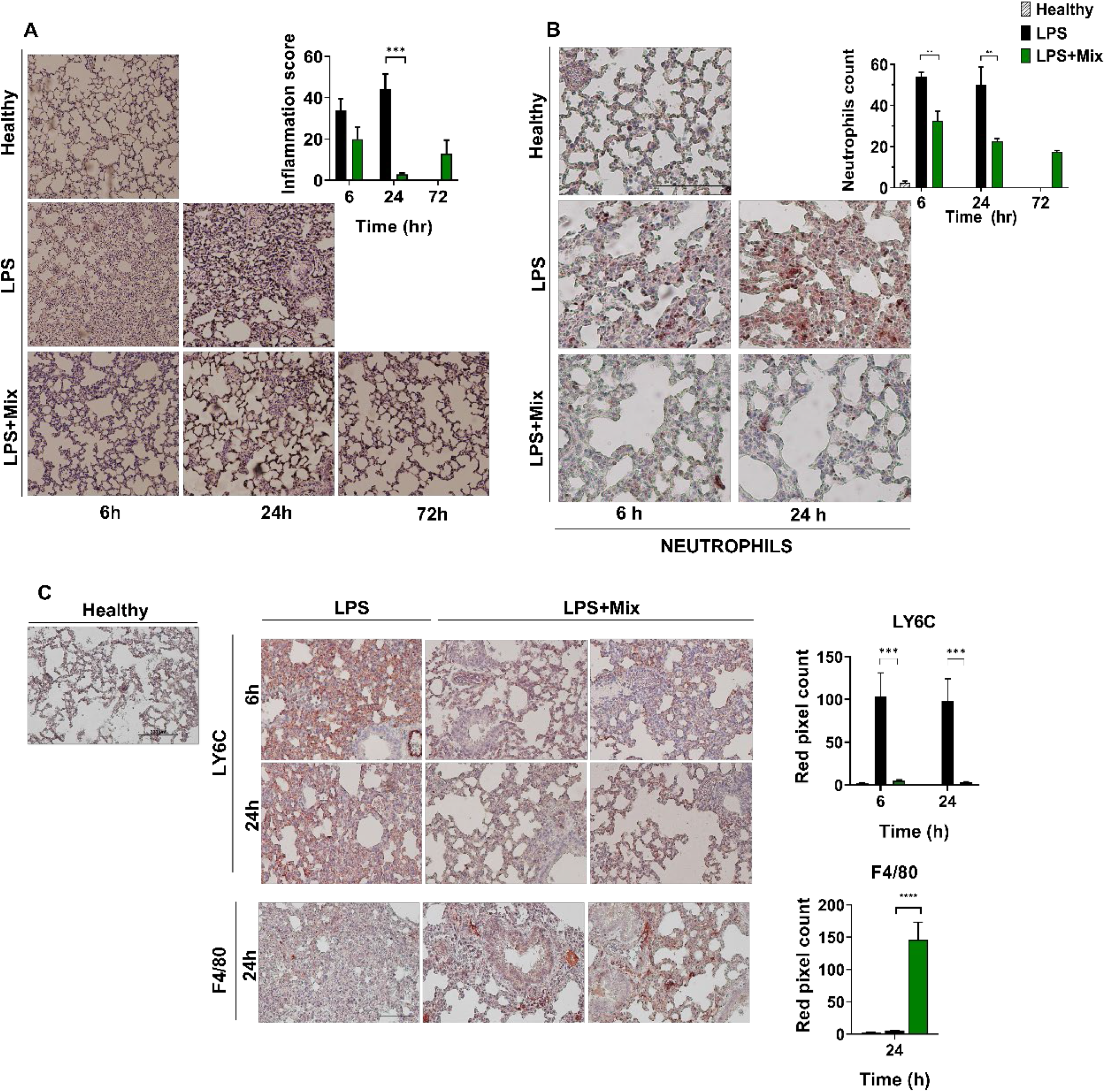
Tryptophol acetate and tyrosol acetate prevent inflammation-associated tissue damage. **A.** Representative pictures of H&E-stained sections of the lung from healthy and LPS-injected mice with or without administration of molecule mixture. Original magnification x20. Quantification of tissue damage was performed, as discussed in Materials and Methods (M&M) and is presented as a bar diagram on top of each panel. **B.** Lung tissues obtained 6 and 24 h after LPS administration were stained with anti-myeloperoxidase (MPO) antibodies for neutrophils detection. Original magnification x10 of the ammount of neutrophils was counts as discribed in M&M and presented in the graph. **C.** Monocytes were detected by anti-Ly6C antibodies and macrophages by anti-F4/80 antibodies. Quantification of the positive cells was assessed, as described in M&M. **p < 0.01; ***p < 0.0001 ****p<0.00001

### Oral uptake of tryptophol acetate and tyrosol acetate decreased tissue damage induced by LPS

Hyperinflammation generally accounts for multisystem failure manifested in considerable damage to the lungs, liver, and the gastrointestinal (GI) tract [29]. To investigate whether the healing effects of tryptophol acetate and tyrosol acetate (e.g., Figure 1) also lead to reduced organ damage, we carried out histology analyses of tissues obtained from LPS-injected mice treated or non-treated with the **1+2** mixture (Figure 2). The representative hematoxylin and eosin (H&E) staining image of lung tissue from a healthy mouse displays normal alveolar walls and no inflammatory cell infiltration (Figure 2A, top row). In LPS-injected mice, signs of severe lung injury characterized by lung edema, hemorrhage, intensive cellular infiltrate, and accumulation of fibrin were observed already during the first 6 hours (Figure 2A, middle panel).

Moreover, lung tissue damage was largely eliminated in mice to which the **1+2** mixture was co-administered with LPS (Figure 2A, bottom row). Indeed, the pulmonary architecture was mostly preserved in those mice, with only small local zones of inflammation. In addition, the alveolar walls were almost not altered, and minimal accumulation of inflammatory cells was observed at the indicated time intervals. Notably, after 72 hours (in which only mice treated with the molecules survived), the lung structure was like healthy mice (Figure 2A(i)). Lung tissue damage was assessed by quantification of pathological score (bar diagram in Figure 2A, inset). Indeed, the increase in mean pathological score after LPS administration was dramatically reduced after treatment with **1** + **2** mixture (at 24 and 72 hours). Those findings demonstrate that tryptophol acetate and tyrosol acetate mixture effectively blocked inflammation-induced damage in lungs.

To further examine the protective effects of the molecules on lung damage after LPS-induced inflammation, we assessed the recruitment of myeloid cells into the lungs (Figure 2A,ii-iii). In the experiments, we performed immunohistochemical (IHC) staining with anti-MPO antibodies for neutrophils, anti-Ly6C antibodies for monocytes (Ly6C^positive^) and anti-F4/80 antibodies for macrophages (F4/80^pos^), considered mature macrophages [30]. Figure 2B, top row, shows minimal neutrophil infiltration in lungs from control mice. However, lungs of LPS-treated mice displayed recruitment of neutrophils already 6 hours after LPS injection (Figure 2B, middle row). Remarkably, in mice which received the **1**+**2** mixture, the recruitment of neutrophils was decreased and was comparable to healthy (PBS treated) mice (bar diagram in Figure 2B, inset). Figure 2C illuminates the recruitment of monocytes and macrophages into the lungs. Both types of cells are strongly associated with inflammation-induced lung pathogenesis [31,32]. Lung-recruited Ly6C^pos^ monocytes account for microvascular endothelial cell activation and vascular injury in LPS-induced early endotoxemia, leading to enhanced pulmonary vascular leakage [33,34]. Figure 2C shows that while the abundance of lung recruited Ly6C^pos^ monocytes increased following LPS administration both after 6 and 24 hours, treatment with tryptophol acetate and tyrosol acetate mixture effectively reduced Ly6C^pos^ monocytes levels back to baseline (i.e. healthy mice). In contrast, treating mice with **1+2** mixture increased F4/80^pos^ myeloid cells (Figure 2C, bottom row), as these macrophages are associated with suppression of inflammation in lung tissues [30,35]. Similar prevention of LPS-induced tissue damage by tryptophol acetate and tyrosol acetate was also apparent in the liver and small intestine (Figure 2, SI).

### Effects of oral uptake of tryptophol acetate and tyrosol acetate on systemic inflammation

Bioavailability analysis indicated that tryptophol acetate and tyrosol acetate appeared in blood 15 min after oral uptake (Figure 3, SI). Accordingly, we further assessed the effects of the molecules on systemic inflammation, as manifested in blood analysis. In the experiments depicted in Figure 3, the peripheral cell blood count (CBC) was evaluated at different time intervals. The results show that in LPS-injected mice that were not treated with **1+2,** reductions in red blood cell (RBC) counts, hemoglobin (HGB) and hematocrits (HCT) occurred. Notably, however, addition of **1** + **2** mixture increased these parameters (Figure 3A, green lines vs black lines). Treating mice with **1** + **2** mixture also increased thrombocyte counts (PLT), which were reduced after LPS injection (Figure 3A). The CBC data in Figure 3A indicate that tryptophol acetate and tyrosol acetate prevented both anemia and thrombocytopenia that are usually associated with severe inflammation, especially accompanied by high secretion of cytokines [36].

**Fig. 3.**
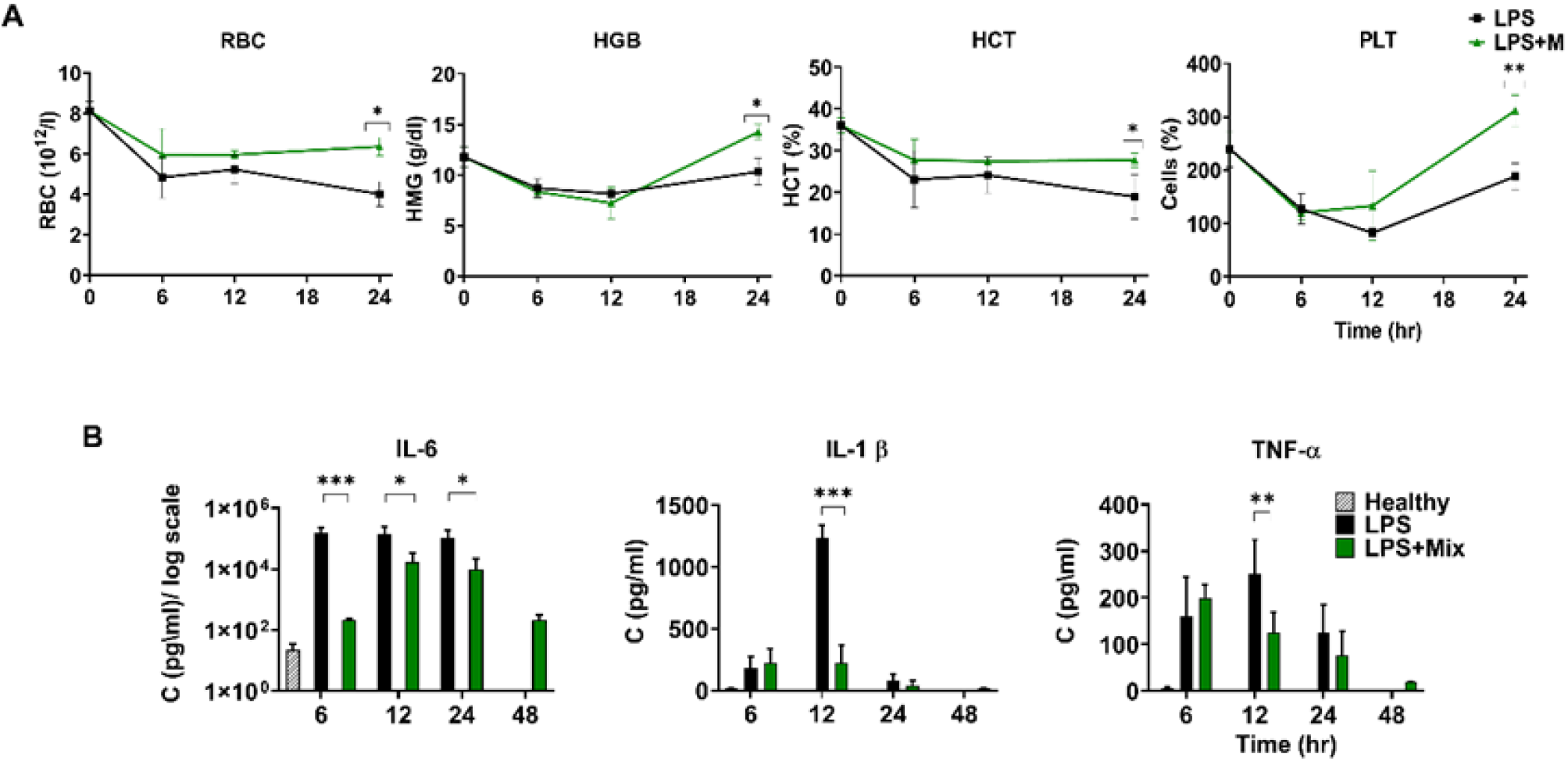
Effects of tryptophol acetate and tyrosol acetate mixture on systemic inflammation. Samples of the peripheral blood were obtained from the tail vein at different time intervals and assessed for CBC. Serums were collected and studied by ELISAs. **A**. The results of RBC, HGB, HCT, and PLT at different time intervals. LPS-administered mice with PBS treatment (black lines) and treated with **1**+**2 mixture** (green), *p < 0.05. **B.** Cytokine levels in serum, assessed by commercial ELISA kits. Data are presented as mean ± SD in each group (n=3). *p<0.05l, **p<0.005, ***p<0.0001.

The clinical hallmark of hyperinflammation is the increased secretion of pro-inflammatory cytokines. Accordingly, we determined the effects of tryptophol acetate and tyrosol acetate mixture given orally to the LPS-injected mice on the levels of pro-inflammatory cytokines, including IL-6, IL-1β and TNF-α in serums (Figure 3B) and lungs (Figure 4, SI). Figure 3B demonstrates that while LPS injection increased the levels of cytokines in serum, treatment with **1**+**2** mixture attenuated cytokine concentrations within 6-24 hours. Specifically, blood concentrations of IL-6 in LPS-administered mice treated with the molecules were lower compared to the untreated LPS-injected mice in all time intervals (note that after 48 hours, LPS-administered mice treated with PBS did not survive, Figure 3B). However, after 48 hours, the IL-6 level of treated mice was almost identical to the healthy mice. Statistically meaningful reductions of IL-1β and TNF-α levels in LPS-administered mice that received the **1+2** mixture were observed 12 hours after LPS injection (Figure 3B). Tryptophol acetate and tyrosol acetate mixture also downregulated mRNA levels of IL-6, IL-1α, and IL-1β in lung tissue at the examined time intervals (Figure 4, SI).

**Fig. 4.**
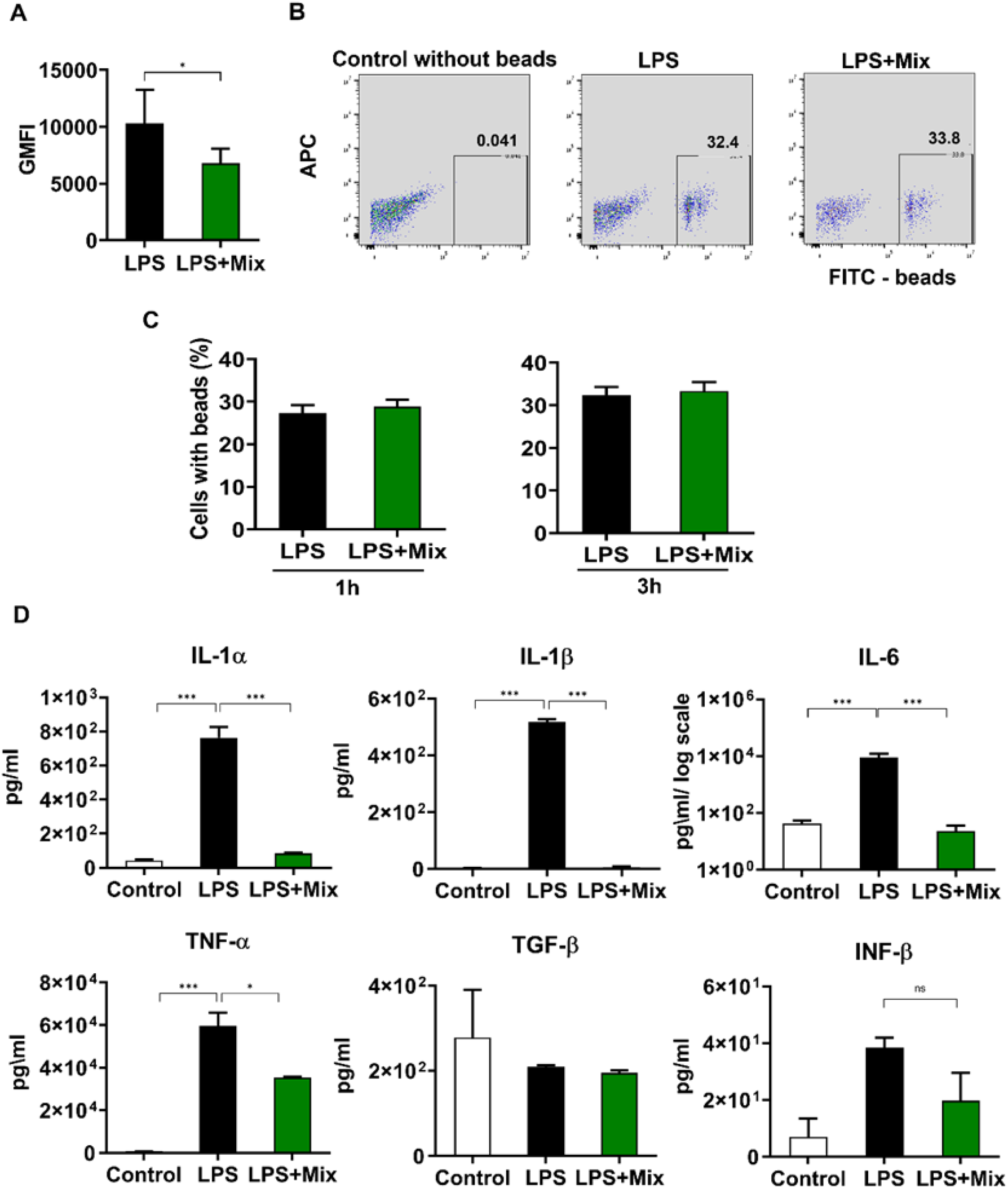
Anti-inflammatory properties of tryptophol acetate and tyrosol acetate *ex vivo*. **A**. Intracellular reactive oxygen species (ROS) after LPS stimulation of murine peritoneal macrophages (three replicates flow cytometry analysis), the results are the Geometric Mean Fluorescent Intensity (GMFI) of the CellRox Deep-Red dye. (Unpaired t-test, p<0.05; n=14). **B**. Representative scatter plots and gating conditions for green-fluorescent beads engulfment analysis, depicting bead uptake after 1 hour incubation of the macrophages, with or without co-addition of **1**+**2** mixture. Percentages of bead positive cells are indicated in the bottom right quadrants. **C**. Analysis of phagocytosis function of murine peritoneal macrophages after LPS stimulation incubated for 1 or 3 hours with or without molecules in concentration of 100 μM each by using green-fluorescent beads (Unpaired t-test, two-tailed; p<0.05). **D**. Cytokine levels determined by ELISA from supernatants of *ex-vivo* isolated peritoneal murine macrophages. Macrophages were incubated for 16-hour with and without LPS and **1**+**2** mixture. Results presented as mean ± SD; n=3 from two independent experiments. *p<0.05 ***p<0.0001.

### Tryptophol acetate and tyrosol acetate reduced the production of ROS and pro-inflammatory cytokines in LPS-activated murine peritoneal macrophages, without reducing phagocytosis function

To further investigate the clinical and biological immunomodulation effects of tryptophol acetate and tyrosol acetate (e.g., Figures 1–3), we performed *ex-vivo* experiments using murine peritoneal macrophages. Figure 4A depicts the effects of tryptophol acetate and tyrosol acetate on reactive oxygen species (ROS) generation using the CellRox Deep-Red dye assay. In general, elevated inflammation levels go together with enhanced production of ROS [37]. Oxidative stress conditions were attained in the experiments by stimulating the murine peritoneal macrophages with LPS (100 ng/ml) with or without co-addition of the **1**+**2** mixture. The flow cytometry analysis revealed an experimentally significant decrease in intracellular ROS after 16 hours incubation of LPS-stimulated macrophages with **1** + **2** mixture, in comparison with the untreated LPS-stimulated macrophages (Figure 4A).

Since phagocytosis is a key function of macrophages [38], we examined macrophage phagocytic action in LPS-stimulated murine peritoneal macrophages with and without co-incubation with the **1** + **2** mixture (Figure 4B,C). Phagocytosis was assessed in the experiments by flow cytometry measuring the uptake of green-fluorescent beads by the cells. The representative gating analysis by flow cytometry (1 hour phagocytosis assay, Figure 4B) together with the bar diagram (Figure 4C) demonstrate no difference in the phagocytic activity of LPS-stimulated macrophages in the absence or presence of the **1**+**2** mixture (1 and 3 hours of phagocytosis assays). These results indicate that the tryptophol acetate and tyrosol acetate mixture did not adversely affect the phagocytic ability of macrophages.

To further confirm the immunomodulatory effects of tryptophol acetate and tyrosol acetate *ex vivo*, we quantified production of pro-inflammatory cytokines by the LPS-stimulated murine peritoneal macrophages with and without co-incubation with the molecule mixture (Figure 4D). Indeed, while the production of IL-1α, IL-1β, IL-6, and TNF-α by macrophages increased within 16 hours after LPS stimulation, addition of the **1**+**2** mixture reduced the concentrations of all these pro-inflammatory cytokines, in most cases back to the non-stimulated macrophage levels. In contrast, tryptophol acetate and tyrosol acetate did not affect secretion of the anti-inflammatory cytokines TGF-β and INF-β (Figure 4D). These results again demonstrate that the addition of the molecule mixture did not disrupt the core anti-inflammatory processes of the host. Together, the results presented in Figure 4 demonstrate that tryptophol acetate and tyrosol acetate decreased intracellular ROS and attenuated generation of pro-inflammatory cytokines, while not adversely affecting important immune processes, including macrophage phagocytic function and anti-inflammatory cytokine production.

### Effect of tryptophol acetate and tyrosol acetate on NF-κB pathways in LPS-activated macrophages ex vivo and in vitro

To obtain insight into the immune modulating activities of the tryptophol acetate and tyrosol acetate mixture, we investigated intracellular pathways associated with NF-κB, a key protein in major inflammation gene cascades [39] (Figure 5). Indeed, the representative western blot (WB) image in Figure 5A (**i**) demonstrates that addition of **1** + **2** mixture to the LPS-stimulated macrophages markedly repressed generation of the phosphorylated form of NF-κB (p65) in comparison to stimulation with LPS (relative band intensities are depicted in the bar diagram in Figure 5A (ii)).

**Fig. 5.**
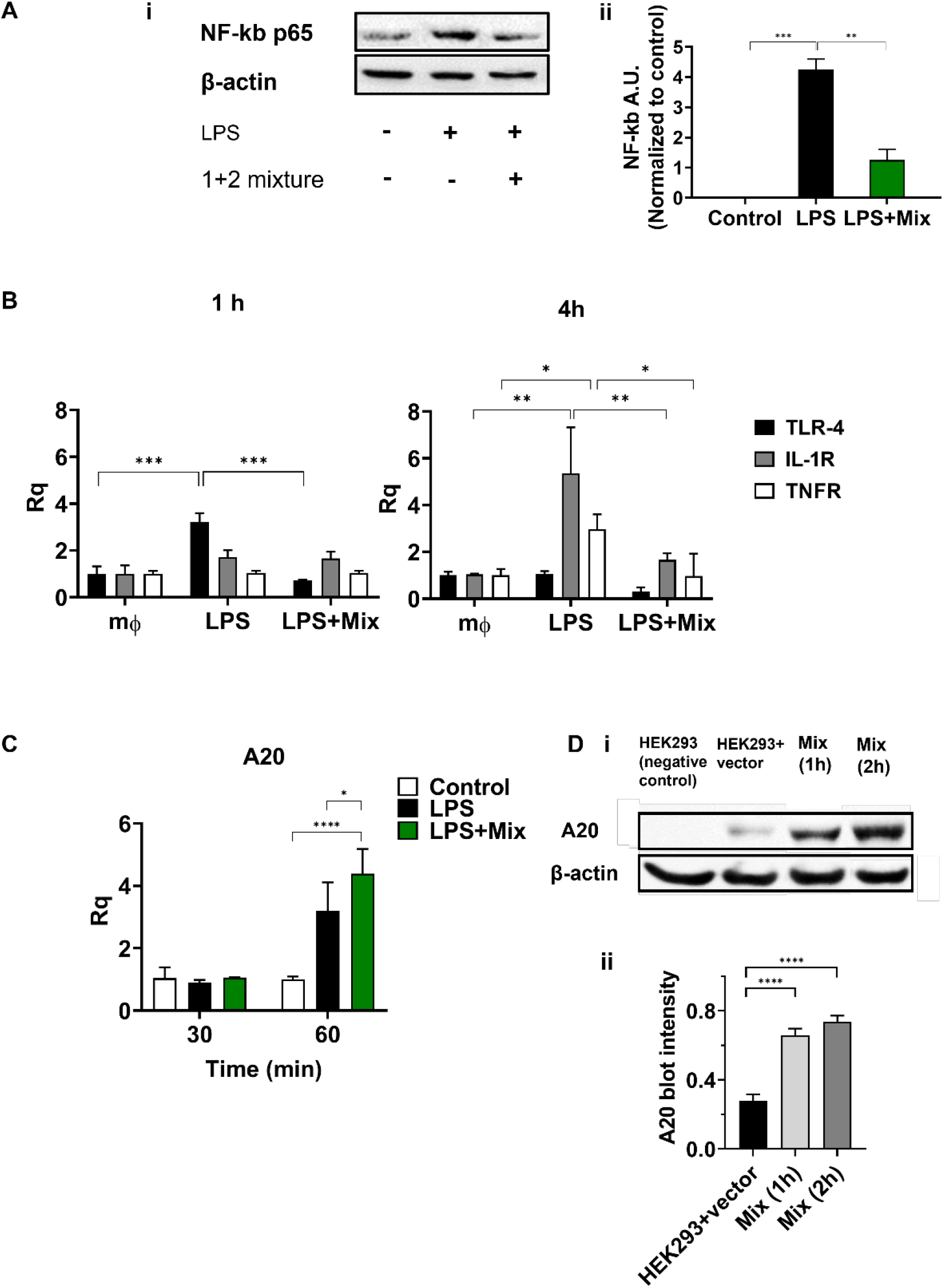
Molecular profiles underlining the anti-inflammatory activity of the tryptophol acetate and tyrosol acetate mixture. Intraperitoneal macrophages were obtained 72 hours after thioglycolate injection and cultured with LPS with or without **1+2** mixture. (**A**) At different time intervals, cell lysates were assessed by Western blot (WB). (**i**) Representative WB of phosphorylated NF-κB (p65) protein expression, with the effects of the mixture (100 μM each). (**ii**) Relative band intensity histograms. The graph represents an average of three independent experiments. Bars indicate standard errors of the means; **p<0.001, ***p<0.0001. Beta-actin was used to assess equal protein loading. (**B**) RAW 264.7 cell line was used for definition of the NF-kB pathway related receptors. Cells were incubated with LPS with and without molecules at the indicated time intervals. RNA was extracted and cDNA was analyzed by RT-PCR. (**C**) Relative protein and mRNA expression of A20 after stimulation with 1 μg/ml of LPS in the absence and presence of **1**+**2** mixture (100μM), detected by RT-qPCR. Error bars indicate standard deviations of three independent cultures. * P<0.05, **** P< 0.0001 ANOVA followed by Tukey’s post hoc analysis. (**D**) (**i**) HEK-293T cells were transfected with pCAGGA vector (*HA-A20*) (1.5 μg) and incubated with and without **1+2** mixture (1 or 2 h). Protein expression of A20 in cell lysates is presented in WB. HEK-293T cells were used as a negative control. (**ii**) Relative band intensity histograms. The graph represents an average of three independent experiments. Bars indicate standard error of the means; statistical analysis calculated by Unpaired t-test, p<0.0001. Beta-actin was used to ascertain equal protein loadings.

To shed further light on the parameters responsible for NF-κB suppression incurred by tryptophol acetate and tyrosol acetate in LPS-stimulated macrophages, we evaluated expression of prominent upstream receptors associated with NF-κB cascades, specifically TLR4, IL-1R and TNFR (40) (Figure 5B). The bar diagrams in Figure 5B depict the levels of the receptors’ mRNA recorded in RAW 264.7 macrophage cells, stimulated by LPS in the presence or absence of **1** + **2** mixture. Figure 5B demonstrates that addition of the molecules to the LPS-stimulated cells significantly attenuated expression of the genes of all three receptors. Specifically, LPS stimulation induced expression of *TLR4* during the first hour (black bar, Figure 5B, left), while co-addition of **1**+**2** mixture together with LPS resulted in suppression of *TLR4* expression, reverting to the baseline (pre-inflammation) level. Similarly, tryptophol acetate and tyrosol acetate significantly attenuated expression of both *IL-1R* (grey bars) and *TNFR* (white bars) 4 hours after LPS stimulation, giving rise to baseline expression levels in both receptors.

Accounting for the reduction of the upstream receptors in the NF-κB pathways by the tryptophol acetate and tyrosol acetate (e.g., Figure 5B), we further tested the effects of the molecules upon expression of the A20 protein, known to suppress signaling cascades associated with the TLR receptors [41] (Figure 5C-D). The bar diagram in Figure 5C presents experimental data corresponding to RAW 264.7 cells treated with **1+2** mixture and the A20 m-RNA levels measured after 30 min and 60 min. As shown in Figure 5C, both LPS, and the **1+2** mixture co-added with LPS, induced the expression of A20 gene within 60 minutes after addition to the cells. However, when **1+2** were co-added to the cells together with LPS, significantly higher expression of A20 was found.

WB analysis was carried out to assess the protein level of A20 (Figure 5D), employing HEK-293T cells transfected with the pCAGGA mammalian expression vector (HA-A20), allowing transient transfection of A20 [22,42]. Specifically, after transfection, the tryptophol acetate and tyrosol acetate mixture was incubated with the cells for different time intervals (HEK-293T cells were used as a negative control, Figure 5D,i). The WB data in Figure 5D demonstrate that **1+2** mixture induced expression of A20, both after 1 and 2 h of incubation compared to the control HEK-293T cells comprising the vector. Together, the RT-PCR and WB results in Figure 5C-D suggest that modulation of A20 levels by tryptophol acetate and tyrosol acetate is likely a key factor in attenuation NF-κB expression and concomitant anti-inflammatory effects of the molecules.

### Tryptophol acetate and tyrosol acetate affect mouse gut microbiome

The relationship between gut microbiota and host immune properties has recently emerged as a prominent factor in systemic response to inflammation. Previous seminal studies, for example, linked inflammatory cytokine production and individual variations in cytokine response to the composition and function of the microbiota [43]. Accordingly, we also examined the effect of tryptophol acetate and tyrosol acetate on the gut microbiome of the mice (Figure 6). In the experiments depicted in Figure 6, mice microbiome analysis was carried out through collecting the stool samples from all participating mice, the day before and at intervals of 6 h and 24 h after LPS injection. The microbial taxonomic 16S rRNA gene sequences were subsequently determined.

**Fig. 6.**
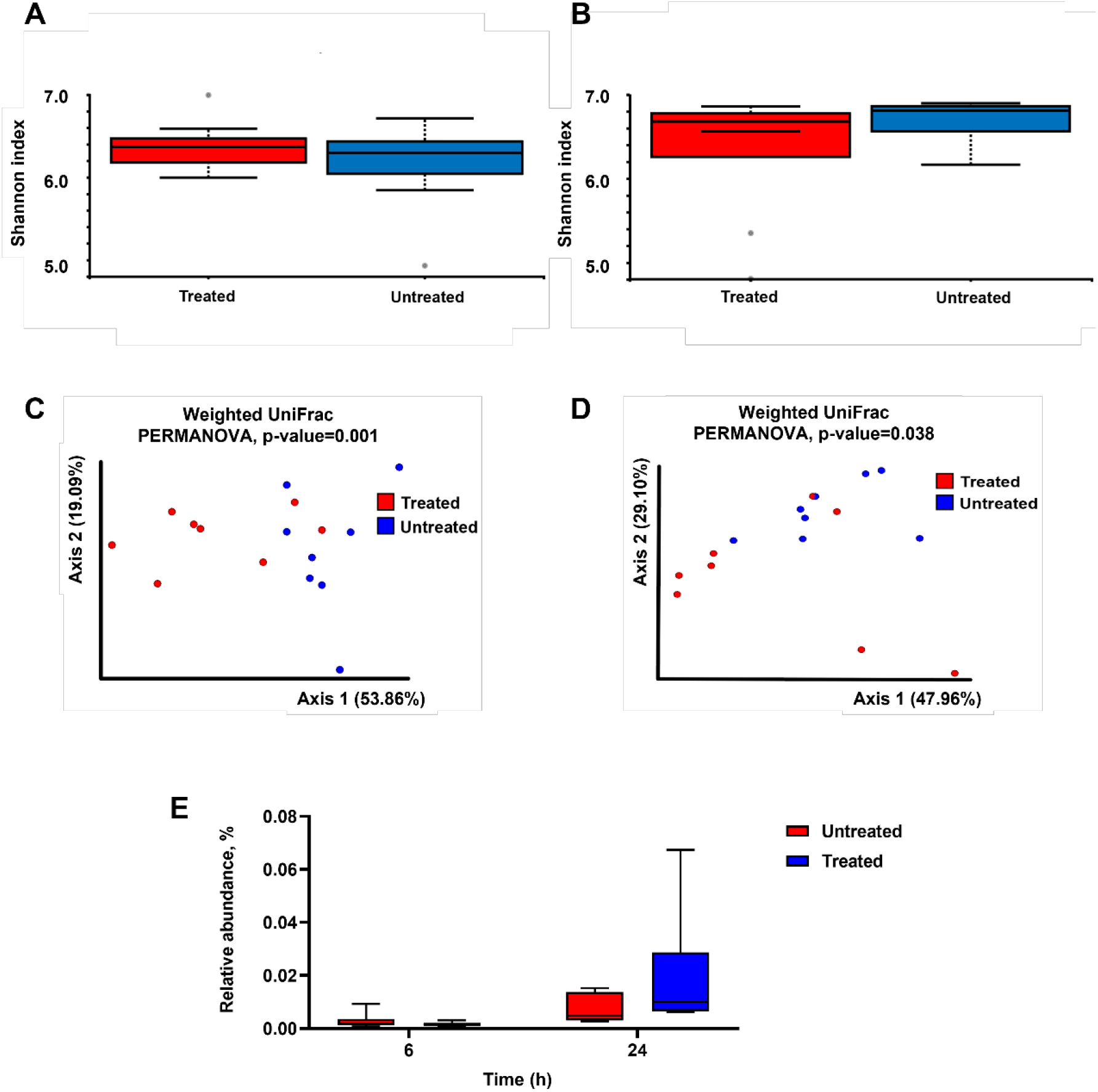
Effect of tryptophol acetate and tyrosol acetate mixture treatment on mice microbiome. **(A, B)** Alpha diversity analysis using the Shannon index after (**A**) 6 hours (p=0.46) and (**B**) 24 hours of treatment (p=0.29); quantitative variables were compared between groups with the Wilcoxon U-test, for non-normally distributed data. **(C, D)** Principal coordinate analysis plot representing beta-diversity based on weighted UniFrac distances after **(C)** 6 hours (permutational multivariate analysis of variance, P=0.001) and **(D)** 24 hours of treatment (permutational multivariate analysis of variance, P=0.038). (**E**) Bacterial taxa *Bacteroides ovatus* relative abundance (%) boxplots for untreated and molecules treated groups. The taxa more abundant in treated group 24 h from LPS injection. (Treated and untreated, n=8)

When comparing the microbial richness (i.e., alpha diversity; Figure 6A, B) before and after treatment with tryptophol acetate and tyrosol acetate (at 6 hours and 24 hours, respectively), no experimentally significant differences were observed (Wilcoxon test, P=0.46 and P=0.29). However, we found differences in beta diversity (i.e., between sample diversities; Figure 6C, D) between treated and untreated mice 6 and 24 hours after LPS injection (PERMANOVA, P=0.001 and P=0.038, respectively). This observation indicates that although the species richness was not significantly altered between the treated and untreated mice, dysbiosis (shift in the microbiome) at the community level was already observed 6 hours after LPS administration. Dysbiosis after LPS treatment was characterized by a reduction in the relative abundance of commensals such as *Blautia*, *Bacteroides*, *Clostridium*, *Sutterella* and *Parabacteroides* and by an increase in the relative abundance of *Ruminococcus* (Table S1, Supporting information). These changes occurred in both the treated and untreated groups. 24 hours after LPS administration we observed a decrease in the relative abundance of *Blautia*, *Bacteroides*, *Clostridium* and *Oscillospira* and increase in the relative abundance of *Ruminococcus*.

Figure 6E underscores the effect of treating the LPS-administered mice with tryptophol acetate and tyrosol acetate. Importantly, the abundance of *Bacteroides* increased in the mice that were treated with the two compounds compared to the untreated mice (24 hours after LPS injection; Figure 6E). This taxon was also abundant in the microbiome of healthy mice (ANCOM test significance W=68 and W=72 respectively (Table S1)). This result is notable as *Bacteroides* has been linked to varied immune-protective and anti-pathogenic activities associated with microbiome modifications [44].

## Discussion

While mortality from sepsis significantly decreased in the past century, in many cases of septic shock or hyperinflammatory syndromes, for example associated with the recent SARS-Covid-19 pandemic, mortality is high [3,45]. Strategies for overcoming sepsis syndromes and associated hyperinflammation have been mostly based on blocking cytokines and other pro-inflammatory molecules. Non-steroid anti-inflammatory drugs (NSAIDs) are the most widely used therapeutic vehicles, however they have many side effects, such as gastrointestinal disorders, water retention, renal failure, bronchospasm, and hypersensitivity reactions [46,47]. Because of the adverse side effects of NSAIDs, traditional medicines and natural products have been promoted as potential alternatives to these drugs, although the efficacy and biomolecular basis for their action have been generally lacking.

Varied food-extracted substances have been touted as exhibiting powerful antioxidant and anti-inflammatory properties [9]. However, reports presenting comprehensive clinical, physiological, and phenomenological analyses of the anti-inflammatory activities of metabolites secreted by probiotic microorganisms have been rare. This work demonstrates pronounced systemic anti-inflammatory properties of two small molecule metabolites – tryptophol acetate and tyrosol acetate - identified in the secretions of the probiotic fungus *Kluyveromyces marxianus* in a milk-fermented microorganism mixture (kefir). When examined in LPS-induced hyperinflammation mice model, the two molecules, given in tandem, effectively blocked mortality and morbidity, and inhibited organ damage (Figures 1–2). In sepsis, multiorgan destruction associated with hyperinflammation leads to hepatic and renal failure, respiratory distress, dysfunction of the gastrointestinal system and hematological changes, such as coagulation disorders [48]. As such, the reported experimental data demonstrate that orally administered tryptophol acetate and tyrosol acetate mixture prevented tissue damage (Figure 2) and attenuated hematological changes (Figure 3).

Circulating mediators of inflammation, such as the pro-inflammatory cytokine IL-6, have been identified as inflammation markers [49,50], and direct links between IL-6 concentration and mortality have been reported [51,52]. Accordingly, the orders-of-magnitude attenuation of IL-6 down to the baseline level after treatment with tryptophol acetate and tyrosol acetate (Figure 3B; Figure 4, SI) further attests to the anti-inflammatory systemic effects of the molecules. This interpretation is also reflected in the lowering of the pro-inflammatory cytokines IL-1β and TNF-α levels in serum few hours after administering tryptophol acetate and tyrosol acetate (Figure 3B). Similar pronounced inhibition effects of pro-inflammatory cytokine levels following treatment with the two molecules were also apparent in *ex vivo* analyses (Figure 4).

Notably, different from numerous anti-inflammatory strategies reported in the literature and clinically employed, tryptophol acetate and tyrosol acetate did not induce complete immunosuppression, rather the molecules affected reverting of the pro-inflammatory cytokine levels to baseline (pre-inflammation) values (Figures 3B, 4D). These observations, together with inhibition of hyperinflammation-induced tissue damage in multiple organs (Figure 2 and Figure 2, SI), and lower anemia and thrombocytopenia (Figure 3A), underscore a systemic therapeutic effect of tryptophol acetate and tyrosol acetate. Notably, the relatively rapid bioavailability of the two molecules (Figure 3, SI) furnishes additional support for their systemic impact.

A major question underlying the anti-inflammatory effects of tryptophol acetate and tyrosol acetate concerns their mode of action. We observed that the molecules have an inhibitory effect on NF-κβ concentration. Furthermore, we recorded downregulation of the main upstream receptors involved in NF-κB activation - TLR4, IL-1R and TNFR (Figure 5A-B). These observations are mechanistically important as it is known that regulation of secreted pro-inflammatory cytokines is governed by the TLR/NF-κB signaling pathway, perceived as a major “gateway” cascade in innate immunity, indeed, modulation of NF-κB signaling cascades have been linked to the pathogenesis of severe inflammation that induces lung, liver, and intestinal injuries [44].

Modulation of A20 protein level by tryptophol acetate and tyrosol acetate [40] (Figure 5) is an important observation accounting for the effects of the two molecules on the TLR4/IL-1R/TNFR NF-κB pathway. A20 is a prominent negative feedback regulator of NF-κB signaling. Mice genetically deficient in A20 develop severe inflammation, underscoring the central role of A20 in suppression of NF-kB-dependent inflammation and tissue homeostasis [53]. Indeed, the observation that tryptophol acetate and tyrosol acetate induced, in both gene and protein levels, A20 expression (Figure 5 C-D) suggest that the protection effect furnished against severe inflammation may be mediated by triggering A20 expression and concomitant inhibition of the TLR4/IL-1R/TNFR-NF-κB pathway.

The gut is thought to play an important role in sepsis pathogenesis [54,55]. Indeed, gut microbiota have been shown to enhance host immunity to pathogens, and dysbiosis has been linked to increased susceptibility of severe inflammation [56]. Since tryptophol acetate and tyrosol acetate are secreted by a probiotic yeast in kefir, it is conceivable that the broad-based anti-inflammatory effects may be also related to modulation of the microbiome upon oral consumption of the molecules. Notably, the predominant change recorded was the increase in the genus *Bacteroides* in mice that were treated with the molecules 24 hours after LPS administration (Figure 6E).

Recent studies have reported on the anti-inflammatory properties of *Bacteroides* [58]. The mechanisms of action proposed for these anti-inflammatory activities include inhibition of pathogen colonization [44] and increased mucosal barrier by modifying goblet cells and mucin glycosylation [59]. The higher abundance of *Bacteroides* upon treating the LPS-injected mice with tryptophol acetate and tyrosol acetate may thus account to “cross-talk” between the molecules and host microbiota, promoting intestinal homeostasis. Moreover, the significance of these results is further underlined by a recent study stressing the need for substances that protect gut microbiota such as *Bacteroides* from the collateral damage of widely used antibiotics such as tetracyclines and macrolides [60]. Thereby, consumption of tryptophol acetate and tyrosol acetate in combination with antibiotics may present a new approach for reducing the harmful side effects of antibiotics on host gut microbiome.

In conclusion, we discovered that orally taken tryptophol acetate and tyrosol acetate - metabolites identified in secretions of a probiotic yeast abundant in a milk-fermented microorganism mixture - inhibit severe inflammation effects. Comprehensive *in vivo, ex-vivo*, and *in vitro* data demonstrate the healing and protective effects of the two molecules, including prevention of mortality in mice experiencing LPS-induced hyperinflammation, blocking of organ damage, inhibition of pro-inflammatory cytokine levels, reduction of ROS production, and maintaining healthy peripheral blood cell profile. Particularly important, unlike many anti-inflammatory strategies, tryptophol acetate and tyrosol acetate did not induce immunosuppression, but rather effectively restored baseline levels of pro-inflammatory cytokines, likely accomplished through downregulation of NF-κB levels via increased A20 expression. Overall, the systemic anti-inflammatory effects of tryptophol acetate and tyrosol acetate, naturally present in a food source consumed by humankind for millennia, may open new avenues for anti-inflammatory therapeutics.

## Acknowledgements

We thank Dr Shai Cohen (Cancer and Vascular Biology Research Center, The Rappaport Faculty of Medicine and Research Institute, Technion - Israel) for providing the A20-plasmid, and his help with the A20 experiments.

## Conflict of interest statement

The authors have no conflicts of interest to declare.

## Author contributions

Conceived and designed the experiments O.M., R.M, E.V. and R.J.; experiments and data analysis O.M., R.M., M.B., M.K., D.C., J.S., E.T., E.S., and B.R.; writing— original draft O.M., R.M., E.V. and R.J.; writing - reviewed and edited the manuscript B.R., O.K., and E.V.; All authors read and approved the final manuscript.

## Notes

### Competing Interest Statement

The authors have declared no competing interest.

### Summary of Updates

Title change, Figures modified

