## supplemental figures for "Tryptophol acetate and tyrosol acetate, small molecule metabolites identified in a probiotic mixture, inhibit hyperinflammation"

### Supporting information

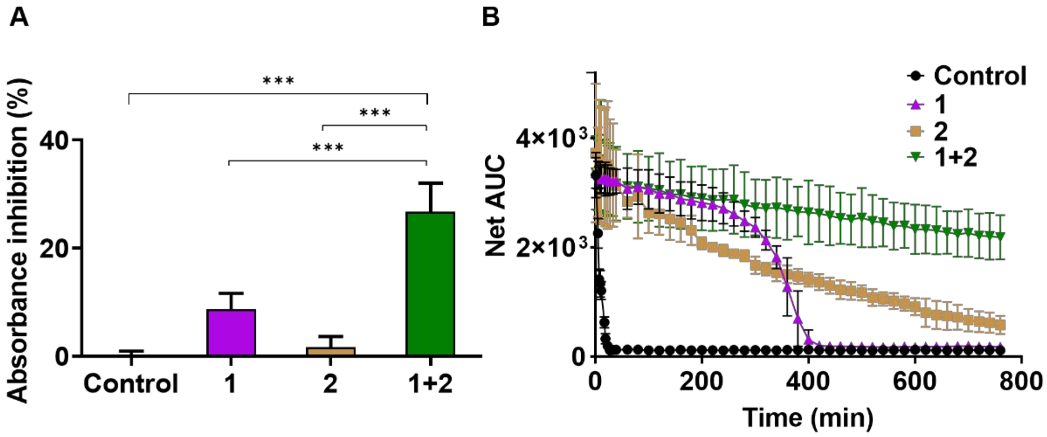


**Fig. 1,SI. Anti-oxidation effect of tryptophol acetate and tyrosol acetate.** Anti-oxidative capacity of the molecules as measured by DPPH (**A**) and ORAC (**B**) assays. **A.** The results are presented as a percentage of absorbance inhibition (517 nm, the absorbance maximum of DPPH) in the presence of tryptophol acetate (**1**) alone, tyrosol acetate alone (**2**) alone, and **1**+**2** together. **B**. Fluorescence decay of fluorescein induced by AAPH in the absence or presence of **1** and **2**. Reaction mixtures containing fluorescein (60 nM) and AAPH (18.75 mM) in 200 µl of phosphate buffer (75 mM, pH 7.4) were incubated at 37°C for 800 min. Changes in fluorescence intensity emitted by fluorescein were monitored. Results are presented as the net area under the curve (AUC). Each value is the mean ± SD of triplicate experiments. ***$p$<0.0001

#### *DPPH scavenging activity*

The effect of tryptophol and tyrosol acetates on DPPH• radical were estimated according to recommendations of Marinova and Batchvarov [52] with some modifications. All solutions were prepared in ethanol. The stock solution was prepared by dissolving 13.8 mg DPPH with 20 mL ethanol and stored until needed. The control (100%) solution was obtained by mixing 225 µL ethanol with 25 µL stock solution to obtain an absorbance of 1.0 ± 0.1 units at 490 nm. 200 µL of tryptophol acetate and tyrosol acetate dissolved in ethanol in a concentration of 200 µM was allowed to react with 25 µL of the DPPH solution for 20 min in the dark at 400 rpm at 25ºC. Ethanol (250 µL) was used for the blank control (100%), and 225 µL was used as a blank. The absorbance decrease was recorded at 490 nm. For all evaluated assays, absorbance measurements were performed in triplicate using a Microtiter Plate Reader (Varioskan Flash, Thermo) to calculate radical scavenging activity (% of inhibition) with the formula.

$$\text{Inhibition (\%)} = 1 - \frac{\text{Abs (sample)} - \text{Abs (blank)}}{\text{Abs (control)} - \text{Abs (blank)}} \times 100$$

where Abs (sample) was the absorbance of the reaction in the presence of the sample (sample dilution + DPPH solution), Abs (blank) was the absorbance of the blank for each sample dilution (sample dilution + DPPH solvent), and Abs (control) was the absorbance of the control reaction (sample solvent + DPPH solution).

***Oxygen radical absorbance capacity (ORAC assay)***

The ORAC method is based on the oxidative degradation of the fluorescent molecules after mixing with a free radical generator, for example, azo compounds. This method determines the ability of the sample to neutralize short-lived free radicals. The assay was carried out according to Ou et al. [53], with some minor modifications. Prior to the measurements, tryptophol acetate and tyrosol acetate were dissolved in acidified methanol to a concentration of 200 µM. 30 µL of each molecule or 30 µL mixture of both, as well as blank (methanol) were mixed with 180 µL of 112 nM fluorescein solution in a 96-well plate and incubated at 37°C for 15 min. Subsequently, 100 µL of 100 mM 2,2' -azobis(2-amidinopropane) dihydrochloride (AAPH) solution was added, and the fluorescence was measured every 70 s for 90 min using a Microtiter Plate Reader (Varioskan Flash, Thermo). The excitation wavelength was 485 nm and the emission recorded at 520 nm. All stock solutions and dilutions of samples were prepared fresh daily in phosphate-buffered saline (PBS, pH 7.4). The experiments were done in six repetitions. Results are presented according to the following calculations :

Area under the curve (AUC) was calculated for each sample using the equation:

$AUC = 1 + FU_1/FU_0 + FU_2/FU_0 + FU_3/FU_0 \ldots\ldots +$

$FU_0$ = fluorescence at time zero.

$FU_x$ = fluorescence at specific timepoints (e.g., $FU_3$ is the fluorescence value at three minutes). Net AUC was calculated by subtracting the Blank AUC from the AUC of each sample using the equation:

Net AUC = AUC (Antioxidant) – AUC (blank)

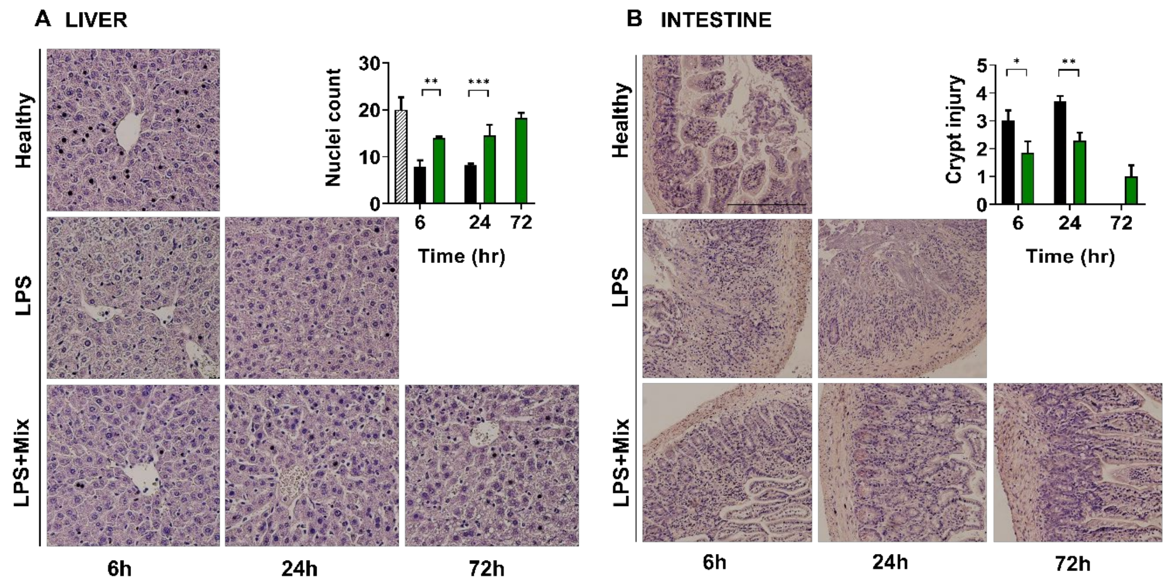


**Fig. 2,SI. Tryptophol acetate and tyrosol acetate prevent inflammation-associated tissue damage. A and B** Representative pictures of H&E-stained sections of obtained organs from healthy and LPS-injected mice with or without administration of molecule mixture **A** liver and **B** small intestine samples. Original magnification x20 for small intestine, x10 for liver. Insert in panel **A** represents an amount of hepatocytes with preserved cell structure. Insert in panel **B** represents quantification of intestinal crypt damage. *$p < 0.05$; **$p < 0.01$; ***$p < 0.0001$

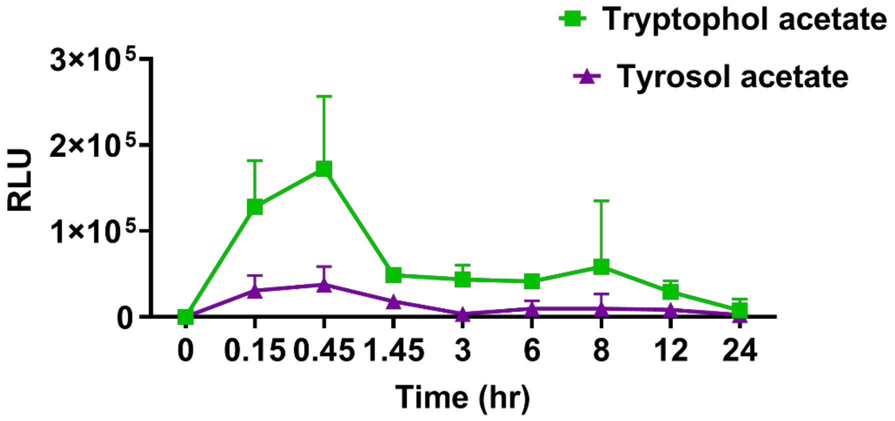


**Fig. 3,SI. Mean levels of tryptophol acetate and tyrosol acetate in serum after oral administration.** n=3 in each time point.

***Bioavailability Assay***

C57BL/6 mice with 20 g body weight were orally dosed by gavage (volume of 200 µL) with 150 µg/kg mixture of tryptophol acetate and tyrosol acetate at 1:1 ratio. Mice (three per time point; male; 10 weeks old) were euthanized with isoflurane at 0.15, 0.45, 1.45, 3, 6, 8, 12, and 24 hours after dosing. Blood was collected by cardiac puncture into syringes and centrifuged for 4 min at 13,000 x g to obtain the serum.

*Sample preparation*: A liquid-liquid extraction was performed by adding 400 µL of ethyl acetate to 100 µM of serum and vertexing the mixture for 3 min. The vortexed samples were centrifuged for 3 min at 10,000 x g, after which each supernatant layer was aspirated and transferred to the GC autosampler vials. Finally, we evaluated the bioavailability of tryptophol acetate and tyrosol acetate by tracking their presence (follow up of characteristic area peaks in the spectra) in the extracted samples using GC-MS as described below.

*Chromatography*:

GC-MS/MS: Agilent; Compatibility: Triple Quadrupole

Column: DB 5 MUIC 0.25 mm × 0.25 mm × 30 m

Gas flow 1 ml/min; Injection volume 4µL

Temperature: QQQ 180°C; MS source 300°C

Software platform: MassHunter

Retention time (RT): tyrosol acetate 11.59 min tryptophol acetate 13.97 min

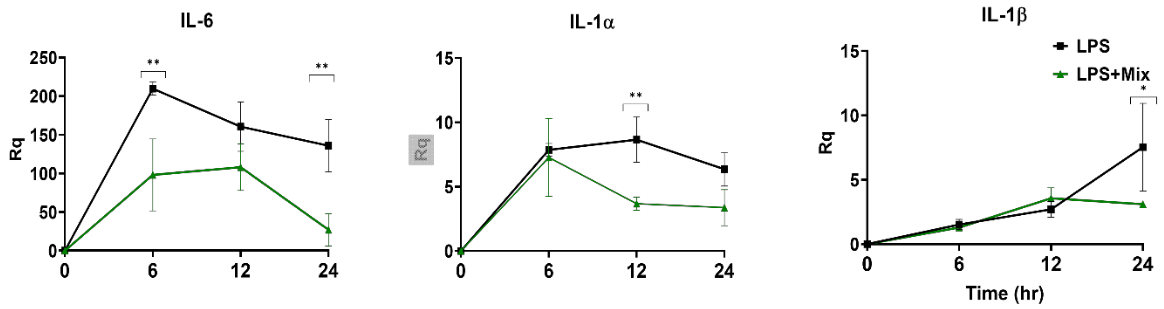


**Fig. 4,SI. Effects of tryptophol acetate and tyrosol acetate mixture on cytokine expression in the lung.** Lungs were obtained from LPS treated mice at indicated time intervals and mRNA was extracted and proceed to RTPCR. Expression of IL-6, IL-1α and IL-1β is shown. Data are shown as mean ± SD in each group (n=4). *p<0.05, **p<0.005.

| ANCOM results after 6 hours | | | |
|---|---|---|---|
| **ASVs** | **Healthy** | **Untreated** | **Treated** |
| *Blautia* | High | Low | Low |
| *Bacteroides* | High | Low | **Low** |
| *Clostridium* | High | Low | Low |
| *Sutterella* | High | Low | Low |
| *Ruminococcus* | Low | High | High |
| *Parabacteroides* | High | Low | Low |

| ANCOM results after 24 hours | | | |
|---|---|---|---|
| **ASVs** | **Healthy** | **Untreated** | **Treated** |
| *Blautia* | High | Low | Low |
| *Bacteroides* | High | Low | **High** |
| *Clostridium* | High | Low | Low |
| *Oscillospira* | Low | High | High |
| *Ruminococcus* | Low | High | High |

**Table 1,SI. ANCOM results of mice treated and untreated with molecules.** Main bacterial taxa abundance after 6 and 24 hours from LPS administration.
